# CameraEEG: Synchronized Recording of Video with Electroencephalogram data on an Android Application

**DOI:** 10.1101/2021.12.21.472324

**Authors:** Srihari Madhavan, Doli Hazarika, Cota Navin Gupta

## Abstract

We present a novel android application named CameraEEG that enables synchronized acquisition of Electroencephalogram(EEG) and camera data using a smartphone. Audio-visual events of interest experienced by the subject were also recorded using a button press on the CameraEEG app. Unlike lab-restricted experiments, which usually constrain the subject’s mobility, this wearable solution enables monitoring of the human brain during everyday life activities. The app was built using Android SDK version 28 and Smarting mobi SDK from mbraintrain. It works on all android devices having a minimum Android OS - Lollipop.

We successfully recorded thirty minutes of synchronized Video and EEG during eyes closed and walking tasks using the app. Event markers enabled by the subject using the app during walking tasks were also recorded. Timing tests showed that temporal synchronization of video and EEG data was good. We analysed the recorded data and were able to identify the task performed by the subject from the event markers. The power spectrum density of the two tasks showed different power spectrums with a peak in the alpha band for eyes closed task. We also provide android studio codes for download and detailed help documentation for the community to test the developed application.

## Introduction

With significant progress in cognitive neuroscience, the current need is to study natural neural responses caused by an ever changing environment in synchrony with the subject’s actions [1]–[3]. In addition, there have been attempts to investigate the changes in neural responses to controlled stimuli in naturalistic experiments [4]. Mobility while measuring brain activity beyond the lab, was made possible by integrating Electroencephalogram (EEG) with smartphones) [5], [6]. Mobile EEG systems have also been used in studying social interaction [7], neural correlates of attention in a college classroom, interpersonal competition and mortality connection through hyper-scanning experiments[8], [9]. Smartphone-based hyperscanning was reported recently to study neural dynamics of interpersonal synchrony between musicians[10]. It is interesting to note that recording camera data with EEG in the above studies might have been useful to locate events of interest.

Clinically, patient monitoring using video-EEG for epilepsy is an accepted practice [11]. Furthermore, mobile EEG systems have been frequently evaluated for epilepsy monitoring and neurodevelopmental disorders [12], [13]. Recently smartphone videos are being studied to diagnosis seizures [14]–[17]. Development and evaluation of a mobile application to record synchronized data from consumer-grade sensors for sleep monitoring was reported in [18]. Overview of smartphone applications for sleep analysis was discussed in [19]. Integrating camera and EEG datastreams definitely seems to be of current interest for epilepsy and sleep diagnosis.

Mobile apps are applications that run on a smartphone for a specific task and recently numerous are being developed for cognitive applications. AFEx and Record-a are two apps developed for understanding sound perception in a real-world situation [2]. Multi-app framework integrated data acquisition (smarting app), BCI processing on SCALA (Signal Processing and CLassification on Android), and stimulus presentation (Stimulus Android app) on an android smartphone [5]. Recently TinnituSense app reported integration of few applications into a single app to record and visualize data in real-time [20]. A variety of sleep analysis apps with their functionality range was presented in [19].

In this work we build a smartphone app named “CameraEEG” which enables synchronous recording of EEG and video. Our aim was to build the app for cognitive experiments in natural environment and hence the back facing camera and EEG data were synchronized. The smartphone was placed in a transparent cell phone hanger to record the video, thereby enabling hands-free mode.

Finally, the system’s performance was tested with a subject doing two tasks namely eyes closed and eyes open. Time lag and continuity between camera and EEG data were checked offline. We analyzed the power spectral density (PSD) of recorded EEG data for both the tasks. To the best of our knowledge we believe our app is the first to implement integration of camera and EEG data on an android system. The code for our app is provided in the following link : https://github.com/Journalsubmissions/CameraEEG

## Methodology

### 1. Software Architecture

Since the application was aimed for an android smartphone, the Android Studio app development software, based on Java Development Kit (JDK), was utilized. Initially, the software utilized the Lab streaming Layer for data acquisition [21] to provide a higher level of time synchronization between different modules and for compatibility with multiple EEG streamer applications. However, this structure was found to be inefficient for our purpose since it required the background streamer app to run in parallel, which drained the battery quicker as well as caused the phone to heat up, causing performance issues. Hence alternatively, the smartphone software development kit (SDK) provided by the EEG manufacturer, specifically the Smarting SDK, was chosen instead. This restricted the use of the app to a specific EEG device (Smarting from mBraintrain) but provided robust execution. This also enabled adding additional modules with unique functionalities to the app.

The app consists of two modules, the camera module, and the EEG input module. The camera module was programmed using the camera2 application programming interface (API) from Android Studio, which provides libraries necessary to capture image frames from the in-built mobile camera unit and stores/access them from a buffer. The camera module contains the video recording and camera preview of the app and is coded based on the ’Camera2Basic’ example by Google (https://github.com/googlearchive/android-Camera2Video), which utilizes a background Handlerthread to parallelize the preview, recording, and other callbacks; and a binary semaphore to prevent opening and closing overlap of the cameradevice object. The resolution of the video recording was set to be 480p (640 × 480) to minimize the storage taken by extended periods of recording (1-2 hrs) and is preset to record from the back-camera due to its higher resolution besides the need to record from the perspective of the subject. The preview window is accessed through a modified SurfaceTexture class named ‘AutofitTexture’ to automatically fit the preview feed from camera to the surface by resizing its dimensions.

### 2. App implementation

The application was developed with the architecture mentioned above. The procedure for data recording (shown in **Figure 1**) in the app was as follows: First, the app was paired to the Bluetooth of the Smarting device, followed by setting up the parameters of connection for recording. Then the app unlocks the recording button (shown in **Figure 2**), which upon pressed, records the synchronous data from the EEG device and the camera video feed into a ‘.bdf’ and ’.mp4’ files, respectively, following which the app can be used to re-record or closed. During the recording, the subject can also press the event marker button to mark any unique event requiring notification.

**Figure 1:**
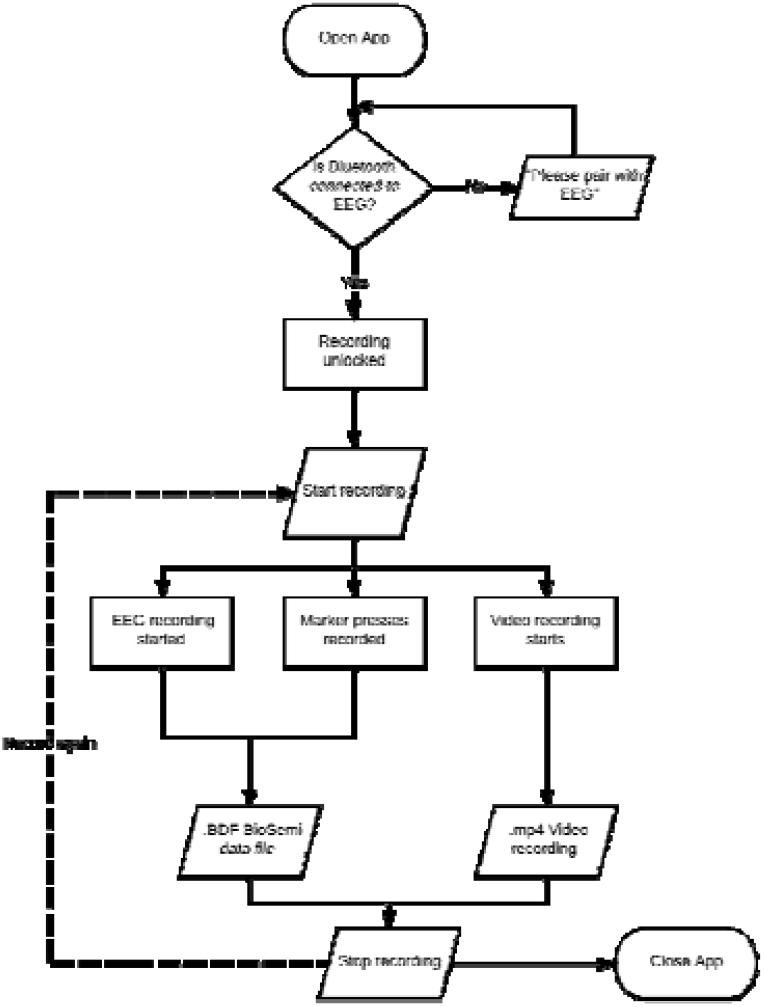
Operation flow diagram of the final application

**Figure 2:**
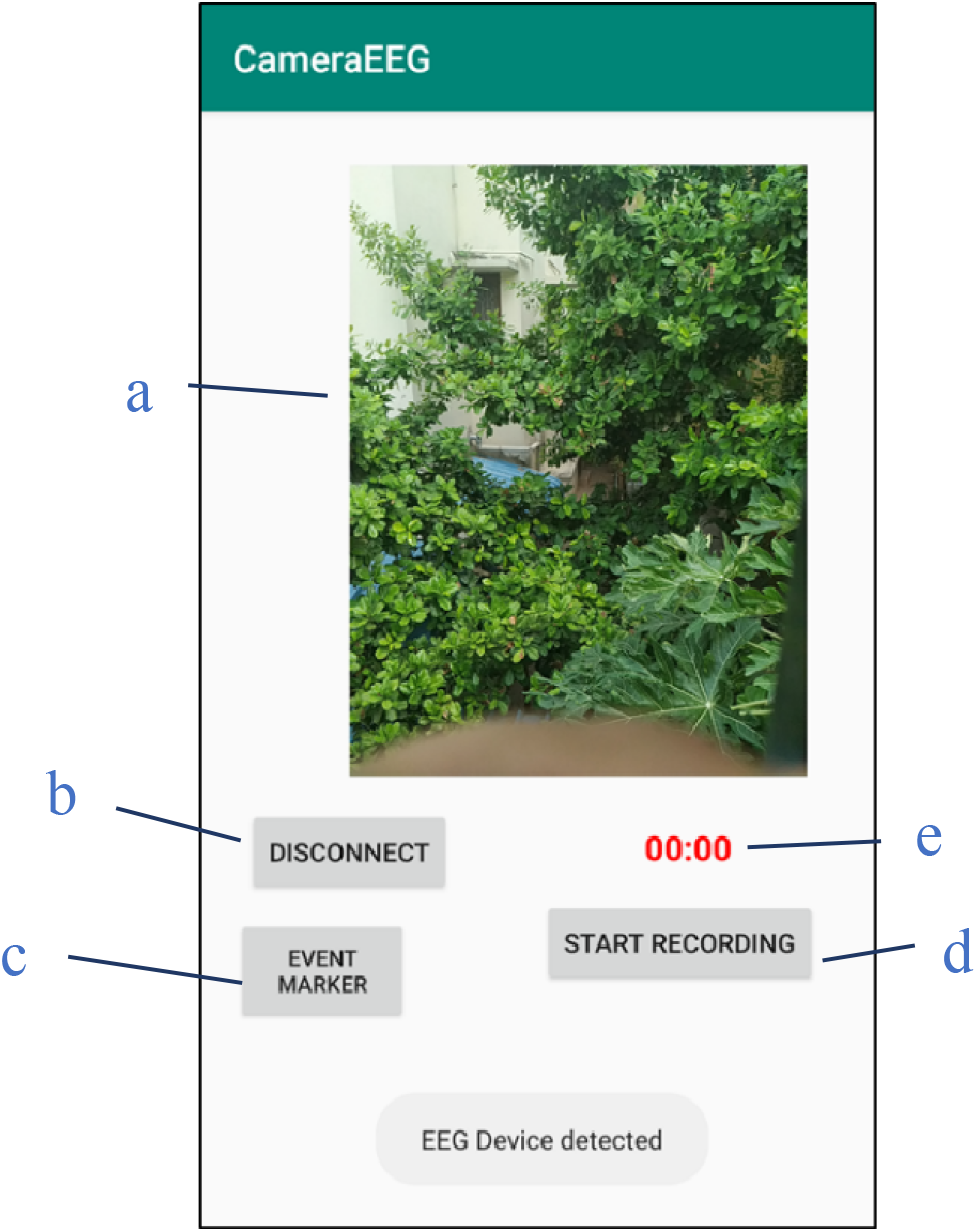
Layout of the app on connecting to EEG device: a) Camera preview window, b) Connect/Disconnect Button, c) Manual event marker button, d) Video-EEG Record Button e) Chronometer-To monitor recording time.

### 3. EEG Recording

Approval was obtained from the Institute human ethics committee, Indian Institute of Technology Guwahati for the recording of data from human subjects. The app was compatible with the 24 channel head-cap of the smarting device, but the 20 channel concealed-EEG (cEEG) was utilized due to its lightweight, mobility, ease of use, and user comfort [22], [23]. It has further been used successfully to record in open, mobile setup [24], study phenomena such as episodic memory[25], selective attention [26], and many more. The smartphone (Sony Xperia Z5 Android 6.1.1) was placed in a transparent plastic cover which allowed video recording through the back-camera while facilitating touch-screen use. As shown in **Figure 3**, It was worn around the neck such that the back camera faced away from the subject, which doesn’t require the cell phone to be held continuously.

**Figure 3.**
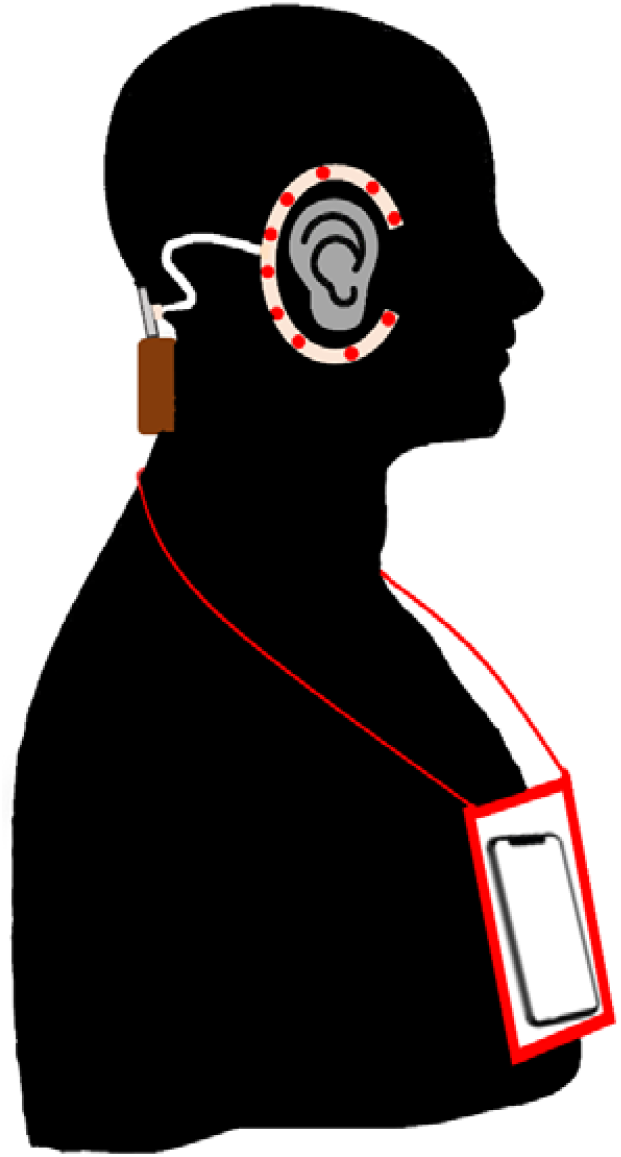
Subject wearing CEEGrid and the app installed android phone

At first the app was tested for data fidelity and synchronization by recording a subject with eyes closed for a short interval of time and while walking for two minutes. This was done to check if changes in activity can be noted from video for EEG event analysis. The change in activity was also noted by pressing the marker button, which notes an event marker to pinpoint the change easier. Upon confirmation of its performance, the subject performed eyes open and eyes closed tasks for upto 30 minutes. With each individual task lasting for 5 minutes.

### 4. Data Analysis

Video from the recording was analysed to check for any data loss. Following this, the duration and time stamps of the activities were noted for analysis.

The EEG data analysis was performed using MATLAB (Version 2020a, The Mathworks Inc., Natick, MA, United States), EEGLAB Version 2020.0, and custom scripts. Upon loading the data, the proper channels of the EEG data were selected, then the length of EEG and video data were compared to check for possible data loss/mismatch between the two. The data was then bandpass filtered between 1 Hz and 45 Hz (FIR filter, filter order: 300). The filtered data were partitioned into segments for sitting with eyes closed and eyes open walking. Following which the alpha band power in both segments, which have been found useful as baselines [27], and compare them along with a paired t-test to confirm the variance between the segments.

### Results

Recording sessions were checked for the duration of EEG and video recording, and they were found to match within an error of ±5ms, with no discernible loss of data. The power spectral analysis of the individual activities (shown in **Figure 4**) was able to show a peak at ∼10Hz for eyes closed, signifying a higher alpha power as expected. The alpha power was quantified by measuring the ratio of power in the alpha band to that of the whole spectrum [28]. The mean increase in alpha band powers of the channels between the activities was 0.49 with an average increase in alpha power ratio of 0.21, and a paired t-test showed a significant difference in means (p<0.0001). We reliably depict that the subject’s activity could be mapped and studied using synchronous Video and EEG. The same result was also visualized in the multisession recordings (shown in **Figure 5**), proving the app’s ability to work under more extended periods.

**Figure 4:**
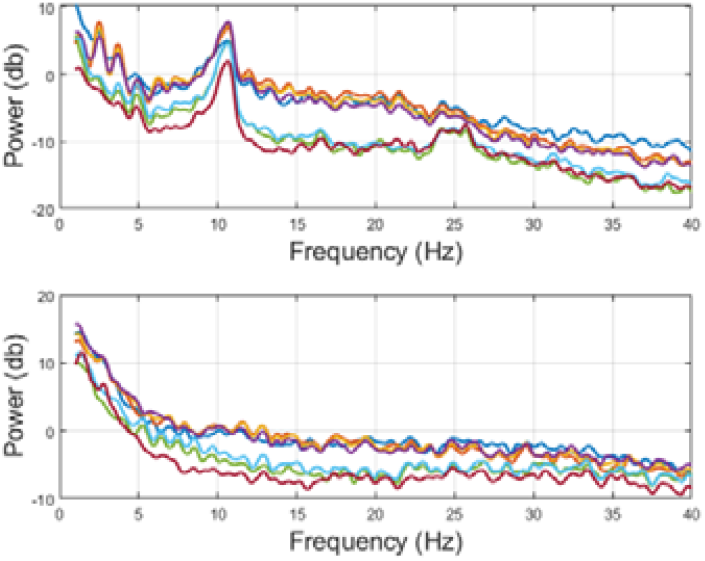
Power spectra of the EEG channels while eyes closed (top) sitting, and eyes open walking (bottom).

**Figure 5.**
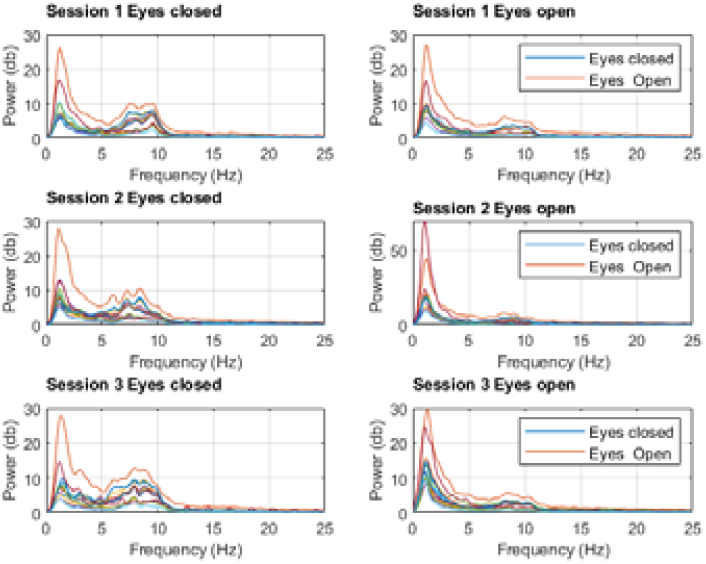
Multisession EEG recordings, eyes open and closed comparisom

## Discussions

The recording sessions of eyes closed and eyes open have been used to validate the performance of our application like in previous works [29], [30]. The peak in the spectra for eyes closed task in comparison to eyes open walking task concur with previous research and shows the proper recording of EEG by the application.

Hence, we show that a good quality video-EEG recording system can be developed using a smartphone. The CameraEEG app provides a simple modular application to monitor and study natural and uncontrolled interactions. Up to our knowledge, no similar system for simultaneous video and EEG recording on a smartphone exists. This system can help record elusive and nuanced social interactions and behaviours, the study of which has gained interest in recent years [31]–[33]. This system can be used as a low-cost epilepsy monitoring, especially in home setups [14], by inclusion of additional functionalities in the developed app.

Unlike previous natural recording applications that seem to use multiple apps or complex setups to present/record external stimuli and EEG [2], [33], our app records them using a single mobile application with EEG device specific libraries. This setup makes it easy to reshape for specific applications and adding on additional modules and decreases the load on the mobile in comparison to using multiple applications simultaneously. However, since our app uses libraries of a particular EEG device manufacturer (https://mbraintrain.com/smarting-mobi/), thereby limiting usage to mbraintain based EEG devices. This is similar to [20] which uses EEG device specific libraries to record and analyse brain data on a smartphone.

While the recording sessions provided primary proof of the app’s functionality, further recordings with multiple subjects and extended durations are required to fully assess the app’s robustness. Older smartphone might heat up a little during the recording sessions but running the app on modern android mobiles (tested on Redmi Note 10 with 4 GB RAM) showed minimal heating. Furthermore, since the recordings were tested only in 480p resolution, recording them in higher resolutions may reduce the performance by heating up the mobile and producing stutters in the app recording. However higher resolutions may provide higher video quality for identifying detailed information about stimuli from the recordings. The current CameraEEG is a stand-alone app that is capable of synchronized recording of EEG and Video.

Further work would be to add a live analysis module which would buffer the live EEG data, apart from writing it to the file, for online analysis such as signal processing, and basic machine learning modules, which could help classify mental states while recording.

## Author Contributions

SM and CNG conceptualized the work, SM designed the application, SM and DH performed data collection and analysis. SM wrote the manuscript with input from DH and CNG.

## Funding sources

SM and DH were funded by Ministry of Human Resource Development scholarship (Master’s and Doctoral scholarship respectively), Government of India. CNG’s time was funded by the Department of Science and Technology (DST), India, and the Swedish Research Council Grant (Project No:14).

## Notes

https://github.com/Journalsubmissions/CameraEEG

### Competing Interest Statement

The authors have declared no competing interest.

## References

[1] S. Georgieva, S. Lester, V. Noreika, M. N. Yilmaz, S. Wass, and V. Leong, “Toward the Understanding of Topographical and Spectral Signatures of Infant Movement Artifacts in Naturalistic EEG,” Front. Neurosci., vol. 14, Apr. 2020, doi: 10.3389/fnins.2020.00352.

[2] D. Hölle, S. Blum, S. Kissner, S. Debener, and M. G. Bleichner, “Real-time audio processing of real-life soundscapes for EEG analysis: ERPs based on natural sound onsets,” bioRxiv, p. 2021.09.20.461034, Sep. 2021, doi: 10.1101/2021.09.20.461034.

[3] M. Jaeger, B. Mirkovic, M. G. Bleichner, and S. Debener, “Decoding the Attended Speaker From EEG Using Adaptive Evaluation Intervals Captures Fluctuations in Attentional Listening,” Front. Neurosci., vol. 14, p. 603, Jun. 2020, doi: 10.3389/FNINS.2020.00603/BIBTEX.

[4] J. Dowsett, M. Dieterich, and P. C. J. Taylor, “Mobile steady-state evoked potential recording: Dissociable neural effects of real-world navigation and visual stimulation,” J. Neurosci. Methods, vol. 332, Feb. 2020, doi: 10.1016/j.jneumeth.2019.108540.

[5] S. Blum, S. Debener, R. Emkes, N. Volkening, S. Fudickar, and M. G. Bleichner, “EEG Recording and Online Signal Processing on Android: A Multiapp Framework for Brain-Computer Interfaces on Smartphone,” Biomed Res. Int., vol. 2017, 2017, doi: 10.1155/2017/3072870.

[6] N. Svenja, J. Jacobsen, S. Blum, K. Witt, S. Debener, and N. Jacobsen, “A walk in the park? Characterizing gait-related artifacts in mobile EEG recordings,” Eur. J. Neurosci, vol. 00, pp. 1–20, 2020, doi: 10.1111/ejn.14965.

[7] F. Babiloni and L. Astolfi, “Social neuroscience and hyperscanning techniques: Past, present and future,” Neurosci. Biobehav. Rev., vol. 44, pp. 76–93, Jul. 2014, doi: 10.1016/J.NEUBIOREV.2012.07.006.

[8] J. K. Grammer, K. Xu, and A. Lenartowicz, “Effects of context on the neural correlates of attention in a college classroom,” npj Sci. Learn. 2021 61, vol. 6, no. 1, pp. 1–4, Jul. 2021, doi: 10.1038/s41539-021-00094-8.

[9] X. Zhou, Y. Pan, R. Zhang, L. Bei, and X. Li, “Mortality threat mitigates interpersonal competition: an EEG-based hyperscanning study,” Soc. Cogn. Affect. Neurosci., vol. 16, no. 6, pp. 621–631, May 2021, doi: 10.1093/SCAN/NSAB033.

[10] A. Zamm, C. Palmer, A.-K. R. Bauer, M. G. Bleichner, A. P. Demos, and S. Debener, “Behavioral and Neural Dynamics of Interpersonal Synchrony Between Performing Musicians: A Wireless EEG Hyperscanning Study,” Front. Hum. Neurosci., vol. 0, p. 476, Sep. 2021, doi: 10.3389/FNHUM.2021.717810.

[11] G. D. Cascino, “Video-EEG Monitoring in Adults,” Epilepsia, vol. 43, no. SUPPL. 3, pp. 80–93, Jun. 2002, doi: 10.1046/j.1528-1157.43.s.3.14.x.

[12] J. Askamp and M. J. A. M. van Putten, “Mobile EEG in epilepsy,” Int. J. Psychophysiol., vol. 91, no. 1, pp. 30–35, Jan. 2014, doi: 10.1016/j.ijpsycho.2013.09.002.

[13] A. Lau-Zhu, M. P. H. Lau, and G. McLoughlin, “Mobile EEG in research on neurodevelopmental disorders: Opportunities and challenges,” Developmental Cognitive Neuroscience, vol. 36. Elsevier Ltd, p. 100635, Apr. 01, 2019, doi: 10.1016/j.dcn.2019.100635.

[14] D. Dash et al., “Can home video facilitate diagnosis of epilepsy type in a developing country?,” Epilepsy Res., vol. 125, pp. 19–23, Sep. 2016, doi: 10.1016/j.eplepsyres.2016.04.004.

[15] B. Ramanujam, D. Dash, and M. Tripathi, “Can home videos made on smartphones complement video-EEG in diagnosing psychogenic nonepileptic seizures?,” Seizure, vol. 62, pp. 95–98, Nov. 2018, doi: 10.1016/j.seizure.2018.10.003.

[16] W. O. Tatum et al., “Video quality using outpatient smartphone videos in epilepsy: Results from the OSmartViE study,” Eur. J. Neurol., vol. 00, p. ene.14744, Feb. 2021, doi: 10.1111/ene.14744.

[17] U. Amin, C. T. Primiani, S. MacIver, A. Rivera-Cruz, A. T. Frontera, and S. R. Benbadis, “Value of smartphone videos for diagnosis of seizures: Everyone owns half an epilepsy monitoring unit,” Epilepsia, vol. 62, no. 9, pp. e135–e139, Sep. 2021, doi: 10.1111/EPI.17001.

[18] A. Burgdorf et al., “The mobile sleep lab app: An open-source framework for mobile sleep assessment based on consumer-grade wearable devices,” Comput. Biol. Med., vol. 103, pp. 8–16, Dec. 2018, doi: 10.1016/j.compbiomed.2018.09.025.

[19] A. A. Ong and M. B. Gillespie, “Overview of smartphone applications for sleep analysis,” World J. Otorhinolaryngol. Neck Surg., vol. 2, no. 1, pp. 45–49, Mar. 2016, doi: 10.1016/J.WJORL.2016.02.001.

[20] M. Mehdi et al., “TinnituSense: A mobile electroencephalography (EEG) smartphone app for tinnitus research,” PervasiveHealth Pervasive Comput. Technol. Healthc., pp. 252–261, Dec. 2020, doi: 10.1145/3448891.3448933.

[21] C. Kothe, “Lab streaming layer (LSL),” https://github.com/sccn/labstreaminglayer. Accessed Oct., 2014.

[22] S. Debener, R. Emkes, M. De Vos, and M. Bleichner, “Unobtrusive ambulatory EEG using a smartphone and flexible printed electrodes around the ear,” Sci. Rep., vol. 5, pp. 1–11, 2015, doi: 10.1038/srep16743.

[23] M. G. Bleichner and S. Debener, “Concealed, unobtrusive ear-centered EEG acquisition: Ceegrids for transparent EEG,” Front. Hum. Neurosci., vol. 11, no. April, pp. 1–14, 2017, doi: 10.3389/fnhum.2017.00163.

[24] S. Debener, F. Minow, R. Emkes, K. Gandras, and M. de Vos, “How about taking a low-cost, small, and wireless EEG for a walk?,” Psychophysiology, vol. 49, no. 11, pp. 1617–1621, 2012, doi: 10.1111/j.1469-8986.2012.01471.x.

[25] M. Piñeyro Salvidegoitia, N. Jacobsen, A. K. R. Bauer, B. Griffiths, S. Hanslmayr, and S. Debener, “Out and about: Subsequent memory effect captured in a natural outdoor environment with smartphone EEG,” Psychophysiology, vol. 56, no. 5, pp. 1–15, 2019, doi: 10.1111/psyp.13331.

[26] W. Nogueira et al., “Decoding selective attention in normal hearing listeners and bilateral cochlear implant users with concealed ear EEG,” Front. Neurosci., vol. 13, no. JUL, pp. 1–15, 2019, doi: 10.3389/fnins.2019.00720.

[27] R. J. Barry, A. R. Clarke, S. J. Johnstone, C. Magee, and J. A. Rushby, “EEG differences between eyes-closed and eyes-open resting conditions,” Clin. Neurophysiol., vol. 118, no. 12, pp. 2765–2773, Dec. 2007, doi: 10.1016/j.clinph.2007.07.028.

[28] G. B. S. Seco, G. J. L. Gerhardt, A. A. Biazotti, A. L. Molan, S. V. Schönwald, and J. L. Rybarczyk-Filho, “EEG alpha rhythm detection on a portable device,” Biomed. Signal Process. Control, vol. 52, pp. 97–102, Jul. 2019, doi: 10.1016/j.bspc.2019.03.014.

[29] A. D. Bateson and A. U. R. Asghar, “Development and Evaluation of a Smartphone-Based Electroencephalography (EEG) System,” IEEE Access, vol. 9, pp. 75650–75667, 2021, doi: 10.1109/ACCESS.2021.3079992.

[30] M. G. Bleichner and R. Emkes, “Building an Ear-EEG System by Hacking a Commercial Neck Speaker and a Commercial EEG Amplifier to Record Brain Activity Beyond the Lab,” J. Open Hardw., vol. 4, no. 1, Oct. 2020, doi: 10.5334/JOH.25.

[31] J. Bhattacharya, “Cognitive Neuroscience: Synchronizing Brains in the Classroom,” Curr. Biol., vol. 27, no. 9, pp. R346–R348, 2017, doi: 10.1016/j.cub.2017.03.071.

[32] G. Cheron et al., “Brain oscillations in sport: Toward EEG biomarkers of performance,” Front. Psychol., vol. 7, no. FEB, 2016, doi: 10.3389/fpsyg.2016.00246.

[33] J. E. M. Scanlon, K. A. Townsend, D. L. Cormier, J. W. P. Kuziek, and K. E. Mathewson, “Taking off the training wheels: Measuring auditory P3 during outdoor cycling using an active wet EEG system,” Brain Res., vol. 1716, pp. 50–61, 2019, doi: 10.1016/j.brainres.2017.12.010.

